# FCGRT, a cancer-derived immunoglobulin G binding protein, mediates the malignant phenotype of glioma

**DOI:** 10.1101/2022.12.23.521719

**Authors:** Guohui Wang, Zheng Wang, Tingting Zhang, Hongyao Ge, Jie Pan, Wangyang Yu, Tianfang Yan, Wei Jiang, Gaoshan Yang

**Affiliations:** Department of Radiation Oncology, Tianjin First Center Hospital, Tianjin, China; Department of Neuro-Oncology, Cancer Center, Beijing Tiantan Hospital, Capital Medical University, Beijing, China; Department of Pathology, Stanford University School of Medicine, Stanford, California, USA; Department of Biochemistry and Molecular Biology, College of Basic Medicine, Hebei University of Chinese Medicine, Shijiazhuang, Hebei, China; Department of Neurosurgery, the Second Hospital of Hebei Medical University, Shijiazhuang, Hebei, China; Department of Neurological Diagnosis and Restoration, Osaka University Graduate School of Medicine, Osaka, Japan; Hebei Higher Education Institute Applied Technology Research Center on TCM Formula Preparation, Shijiazhuang, Hebei, China

**Keywords:** FCGRT, glioma, tumor microenvironment, prognosis, cancer-derived immunoglobulin G

## Abstract

CIgG has received increasing attention, and was first discovered by our group to indicate poor prognosis in glioma. Furthermore, by protein mass spectrometry, we found that FCGRT can combine with CIgG. However, the study of FCGRT in glioma has not been reported. We used the CGGA325, TCGA dataset and immunohistochemistry to verify the importance of FCGRT on the prognosis of glioma patients. Single cell sequencing data analysis evaluated that the role of FCGRT in the microenvironment of glioma. Estimate, ssGSEA, EPIC and xCell were used for immune infiltration analysis. FCGRT was knocked down in U251 cells to detect the effect of FCGRT on the malignant development of glioma. These results showed that patients with higher FCGRT expression had a shorter overall survival. FCGRT was closely related to the tumor microenvironment, especially to macrophages in the tumor microenvironment (r=0.743, p<0.001). Interestingly we also found that FCGRT was positively correlated with IGHG1. Finally, we found that knock-down of FCGRT resulted in a decrease in proliferation, migration and invasion of U251 cells. Taken together, we believe that FCGRT is an independent prognostic factor for glioma patients, and its possible mechanism is to promote proliferation and invasion of tumor cells by interacting with CIgG.

## 1. Introduction

Gliomas, which originate from glia cells, are the most frequently diagnosed intracranial malignancies[1]. Gliomas can be divided into grades 1–4; grades 2–3 are lower grade gliomas, and grade 4 is the highest grade glioma, also known as glioblastoma (GBM). However, the latest WHO classification has introduced molecular features for classifying gliomas, such as IDH, 1p19q and MGMT[2,3]. Treatment of gliomas usually involves surgery, chemotherapy, or radiotherapy. However, the prognosis of patients with gliomas is poor. In particular, patients with GBM, who are given radiotherapy in combination with temozolomide chemotherapy, following complete surgical resection, have a 5-year survival rate of just 6.8%[4]. This is primarily because glioma cells grow rapidly and show infiltrative growth, which widely infiltrates adjacent brain tissue[5]. Immunotherapy is highly efficient in other solid tumors, but the therapeutic effect in glioma is unsatisfactory, possibly due to the tumor microenvironment in the brain being different from other tumors.

The classic immunological theory specifies that immunoglobulin G, produced by B lymphocytes, protects against foreign microbes, exerting the effects of antibodies. However, an increasing number of scholars have observed that tumor cells can also produce immunoglobulin G, and are thus known as cancer-derived immunoglobulin G(CIgG). Several studies have shown that CIgG promotes tumor progression[6–8]. Our group was the first to show that CIgG is strongly associated with a poor prognosis of glioma patients, and may be involved in shaping the immunosuppressive microenvironment of gliomas[9]. In a follow-up study, our group found—through an immunoprecipitation-mass spectrometry study—that CIgG could combine with Fc gamma receptor and transporter (FCGRT). FCGRT encodes a receptor that binds the Fc region of monomeric immunoglobulin G. Binding of FCGRT to immunoglobulin G protects the antibody from degradation[10].

In this study, we verified the relationship between FCGRT and glioma prognosis, and the possible underlying mechanism, by means of biological information and in vitro experiments.

## 2. Materials and Methods

### 2.1 Data acquisition

The CGGA325 database including RNA sequencing and clinical characteristic information was downloaded from the Chinese Glioma Genome Atlas (http://www.cgga.org.cn/)[11]. Similarly, TCGA RNA sequencing data and clinical information were downloaded from the TCGA database (https://portal.gdc.cancer.gov/) as a validation set. Single cell data was downloaded from GSE117891[12], with more than 6000 single-cell transcriptomes of 73 surgery points from 13 gliomas and one brain metastasis patient.

### 2.2 Prognostic analysis

We used CGGA325 as a test set and TCGA as a validation cohort for survival analysis. Glioma patients were divided into high and low groups according to the median value of FCGRT expression. Kaplan–Meier survival analysis was used to analyze the correlation of FCGRT expression on the prognosis of glioma patients. Subsequently, we performed a subgroup analysis of glioma patients by dividing the CGGA325 dataset into 12 subgroups based on age (Age<42, Age≥42), gender (Female, Male), grade (Grade 2, Grade 3, and Grade 4), IDH (Mutant, Wildtype), 1p19q (Codel, Non-codel) as well as MGMT status (Methylated, Un-methylated). Survival analysis was performed in each subgroup to analyze the effect of FCGRT on the prognosis of glioma patients. The TCGA database as a validation set was similarly analyzed.

### 2.3 Immunohistochemistry

The tissue microarray (TMA) for the glioma was obtained from the Shanghai Outdo Biotech Company (Shanghai, China). We used the ab64264 kit from Abcam for immunohistochemical staining, and the detailed staining procedure can be seen in the instructions. It should be noted that the antigen retrieval solution we used was Tris-EDTA, pH = 8.0. Immunohistochemical scoring criteria can be seen in previous publication from our team[13].

### 2.4 Single cell analysis

The single cell RNA-seq data was obtained from GSE117891. The single cell RNA-seq was analyzed using the R package Seurat[14]. Single cell quality control standard refers to the previous articles published by our team[15].The Seurat package’s FindMarkers function was used to identify the cell cluster.

### 2.5 Analysis of immune infiltration and tumor microenvironment

We analyzed the relationship between FCGRT and tumor microenvironment by four immune infiltration analysis methods, including ssGSEA, ESTIMATE, EPIC, and xCell. SsGSEA was used to evaluate 24 types of immune cells that may infiltrate into the tumor immune microenvironment. ESTIMATE was to evaluate infiltration of immune cells and the presence of stromal cells in tumor samples, which generated three results, namely immune score, stromal score, and estimate score. EPIC can analyze the infiltration proportion of eight types of immune cells according to the expression data, including B cells, tumor associated fibroblasts (CAFs), CD4+ T cells, CD8+ T cells, endothelial cells, macrophage cells and NK cells. XCell was used to estimate the abundance scores of 64 immune cell types, including adaptive and innate immune cells, hematopoietic progenitors, epithelial cells, and extracellular matrix cells.

### 2.6 Non-negative matrix factorization (NMF) cluster analysis

In order to further explain the relationship between FCGRT and the immune microenvironment, we used the NMF cluster analysis method. Firstly, we downloaded monocyte-macrophage characteristic genes from previous reports[16] We extracted immune related gene expression quantities from the CGGA database and then performed NMF clustering analysis. The optimal number of clusters was calculated according to the values obtained from cophenetic calculations. We then performed survival analysis on different clusters. Finally, we measured the expression levels of FCGRT and immunoglobulin heavy chain gene 1 (IGHG1), and detected the differences in gene expression levels in different clusters by Wilcoxon rank sum test.

### 2.7 Gene set enrichment analysis (GSEA)

We obtained the GSEA software (version 3.0) from the following website: (http://software.broadinstitute.org/gsea/index.jsp). According to the expression levels of FCGRT, samples were divided into a high FCGRT expression group (≥50%) and a low FCGRT expression group (<50%). Methods can be seen in previous publication from our team[9].

### 2.8 Cell culture

The human glioma cell line U251 was obtained from the American Type Culture Collection (ATCC), and routinely cultured in RPMI 1640 medium containing 10% fetal bovine serum (FBS, Gibco) and 100 U/mL penicillin and 100 μg/mL streptomycin, in a humidified incubator at 37°C with 5% CO2. The medium was changed every 2 d, and cells were passaged when the growth density reached about 80%, about every 3 to 4 d.

### 2.9 Cell proliferation assay

1×10^4^ cells/well were seeded in 96-well plates in RPMI 1640 medium, and the proliferation rate of the U251 cells was measured by the MTT assay. After appropriate treatment, cells were washed with PBS and were replaced with 100 μl serum-free RPMI 1640 containing 10 μl of MTT reagent, and then incubated at 37°C for 4 h. An equal volume of supernatant from each well was transferred to a new 96-well plate. The absorbance at 490 nm was measured using a Multiskan spectrum and the absorbance of MTT reagent at 490 nm was used as a control to corrected absorbance.

### 2.10 Isolation of RNA and real-time PCR

Total RNA was extracted from U251 cells with TRIzol reagent (Qiagen, 10296-028) according to the manufacturer’s instructions. Reverse transcription and qRT-PCR were carried out on ABI 7500 FAST system (Life Technologies) according to the manufacturer’s protocol. β-actin was used as an internal control. All PCR experiments were performed in triplicate. Primer sequences were as follows: FCGRT: 5’-GGG GAA AAG GTC CCT ACA CTC-3’ and 5’-CCT GCT TGA GGT CGA AAT TCA T-3’; β-actin: 5’-GGC TGT ATT CCC CTC CAT CG-3’ and 5’CCA GTT GGT AAC AAT GCC ATG T-3’.

### 2.11 Western blot analysis

Proteins were isolated from cultured U251 cells with lysis buffer and were electrophoresed on 10% SDS-PAGE to separate. SDS-PAGE gels were transferred onto PVDF membranes and were blocked with 5% milk in TTBS for 2 h at 37°C. membranes were incubated with primary antibodies overnight at 4°C. Antibodies used were: anti-FCGRT (rabbit polyclonal, 1:500, 16190-1-AP, Proteintech), anti-β-actin (mouse monoclonal, 1:1000, sc-47778, Santa Cruz). Membranes were then washed and incubated with the corresponding secondary antibodies conjugated with HRP (Cell Signaling Technology). Protein bands were detected by ECL (enhanced chemiluminescence) Fuazon Fx (Vilber Lourmat).

### 2.12 Cell transfection

U251 cells were transfected with small interfering RNAs using Lipofectamine 3000. The sequences of human FCGRT (ID: 2217) specific siRNAs and non-specific siRNA (si-Ctl) were designed. After 24 h of transfection, U251 cells were harvested and lysed for qRT-PCR or western blot analysis. FCGRT siRNA1: 5’-GCTCTTTCTGGAAGCTTTCAA −3’; 5’ - CGGCGAGGAGTTCATGAATTT −3’; siRNA2: 5’ - GGAUCUCUCCUACAGGUAAC −3’; 5’ - UUACCUGUAGGAGAGAUCCGA −3’; siCtl: 5’-UUC UCC GAA CGU GUC ACG UTT-3’; 5’-ACG UGA CAC GUU CGG AGA ATT-3’.

### 2.13 Cell viability and colony formation assay

U251 proliferation was assessed by the CellTiter-Glo assay according to the manufacturer’s protocol. U251 cells, parental or with FCGRT knockdown were plated into 96-well plates with a seeding density of 2000 cells/well. Proliferation was assessed using CellTiter-Glo reagent (Promega). U251 cells were plated in six-well plates with a seeding density of 100 cells/well for the colony formation assay; after 14 days, plates were washed with PBS and stained with Geimsa. Geimsa-stained plates were washed after 30 min, and colonies were counted and analyzed.

### 2.14 Cell migration assays

U251 cells transfected with si-Ctl or si-FCGRT were grown to a confluence of 90%, and the cell monolayer was then scratched using a sterile micropipette tip. After replacing the medium, images at 0 h of the wounded area were captured. Cell migration into the wounded area was recorded again after 12 h. The migration abilities of U251 cells were quantified by measuring the change of scratch regions.

### 2.15 Cell invasion assays

Transwell Matrigel invasion assays were used to assess the invasive abilities of U251 cells, as previously described[17]. Experiments were conducted in triplicate, and migration was expressed as the mean ± SEM per field of total cells counted.

### 2.16 Time-lapse experiments

Time-lapse images of U251 cells transfected with si-Ctl or si-FCGRT were acquired by Live Cell Imaging System. U251 cells maintained at 5% CO2 and 37°C on the inverted microscope at all times. Images were recorded every 30 minutes via a 40× objective. After 120 min, the migration distance of cells was observed and quantified by NIH Image J software.

## 3. Results

### 3.1. Characteristics of patients with gliomas

The CGGA325 dataset contains RNA sequencing data and clinical prognosis information from 325 glioma patients. After we removed patients with incomplete clinical information, 286 glioma patients remained. As the validation set, TCGA contains LGG and GBM datasets. We first combined the two datasets, and then removed patients with incomplete clinical information, and finally 592 glioma patients were included in the follow-up analysis. The GSE117891 database contains a total of 6148 cells, involving 73 regions from 13 patients, which was used for single cell analysis. All detailed information is represented in Table 1. The article flow chart is shown in Supplementary FigureA1.

### 3.2. FCGRT as an independent prognostic factor for glioma patients

To explore the relationship between FCGRT and the prognosis of glioma patients, we utilized the CGGA325 dataset and the TCGA cohort for survival analysis. Glioma patients were divided into high and low expression groups according to the median value of FCGRT expression. The CGGA325 dataset showed that the median survival time was 6.5 years (3.1–9.5) in the low FCGRT expression group, and 1.3 years (1– 1.8) in the high FCGRT expression group according to the log rank test (HR=2.69, 95% CI:2.00–3.60, P < 0.001) (Figure 1A). Similarly, in the TCGA database, the median survival time of the low FCGRT expression group was 7.8 years (6.2–11), and of the high FCGRT expression group was 2.2 years (1.6–3). Survival analysis also showed that FCGRT had a strong impact on the prognosis of glioma patients (HR=3.45, 95% CI:2.47–4.83, P < 0.001) (Supplementary FigureA2). According to the clinical characteristics of glioma patients, gliomas are divided into age (Age<42, Age≥42), gender (Female, Male), grade (Grade 2, Grade 3, and Grade 4), IDH (Mutant, Wildtype), 1p19q (Codel, Non-codel) as well as MGMT status (Methylated, Unmethylated). In the CGGA325 dataset, except for Grade 4, the other groups showed that FCGRT could lead to poor prognosis of glioma patients, which was statistically significant (Figure 1B-M). It is particularly interesting to note that FCGRT expression in patients was low in both the 1p19q codel group. It is well established that 1p19q codel is a predictor of better prognosis in glioma patients. It also further illustrated that FCGRT expression was closely related to poor prognosis of patients. Similar results were obtained in TCGA (Supplementary FigureA2). All results were statistically significant except in Grade 3, Grade 4, IDH mutant and 1p19q code groups. The tissue microarray included 87 gliomas and three adjacent noncancerous tissues. According to the score of immunohistochemical staining, low FCGRT expression was detected in three cases of normal brain tissues, high FCGRT expression was detected in 47 (47/87) glioma patients, and low FCGRT expression was detected in 40 (40/87) gliomas. Most FCGRT positive areas were in the cytoplasm and membrane, and there was also a minor distribution in the nuclear envelope. Figure 2A-C shows the low, medium and high expression of FCGRT in glioma. Of the 87 patients with gliomas, 15 had prognostic survival information. We performed survival analysis on these 15 cases. Survival analysis revealed that the survival time was significantly shorter in the high FCGRT expression group than in the low FCGRT expression group (HR=6.41,95% CI:1.31–31.45, P < 0.001) (Figure 2D).

**Figure 1.**
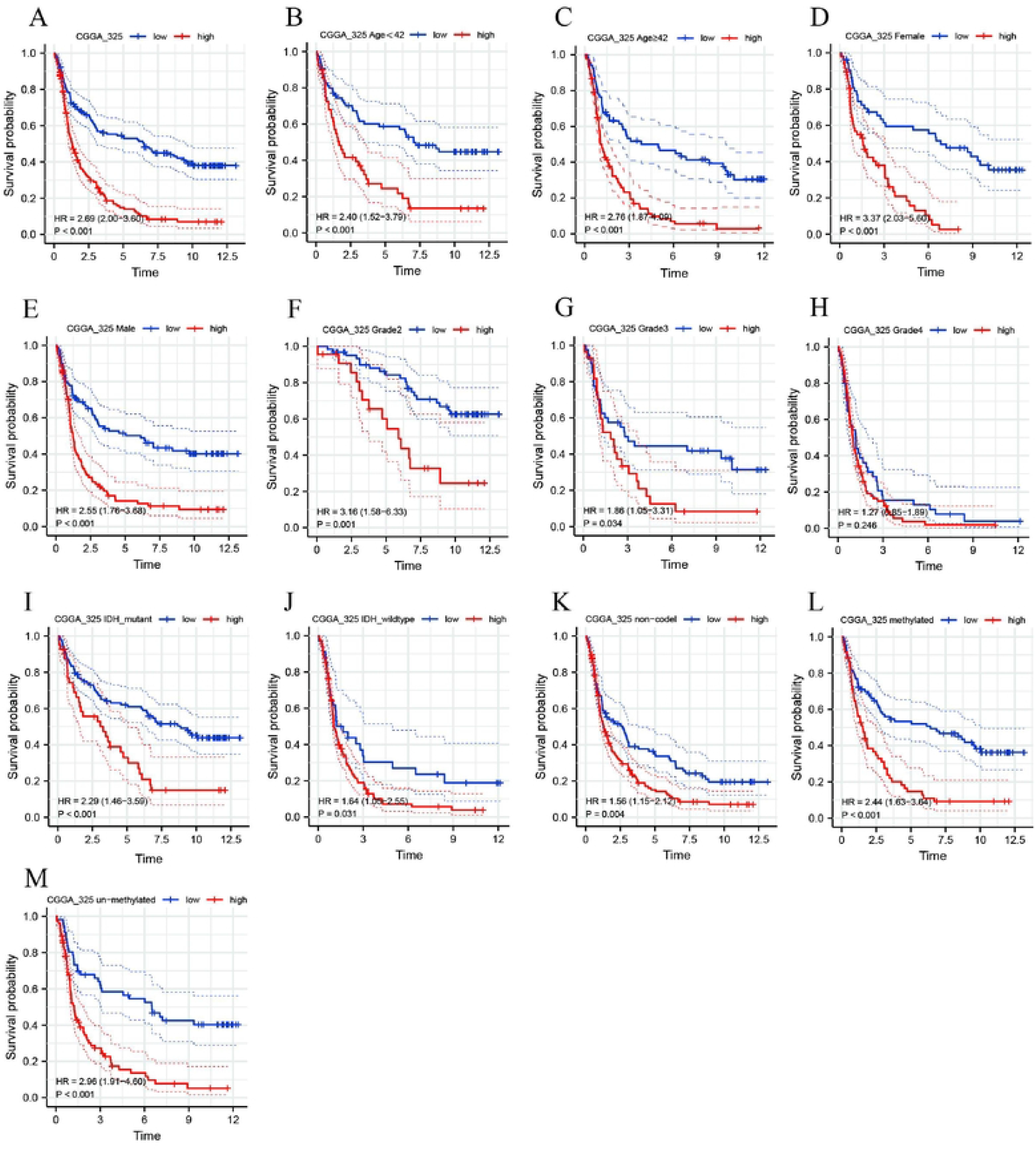
Prognostic analysis of the FCGRT in CGGA325 set. Survival curve was used to analyze OS of the low- and high-risk groups in CGGA325 dataset (A). (B-M) Survival analysis of the FCGRT in patients stratified by age, sex, PRS type, grade, IDH, lp!9q status, and MGMT promoter.

**Figure 2.**
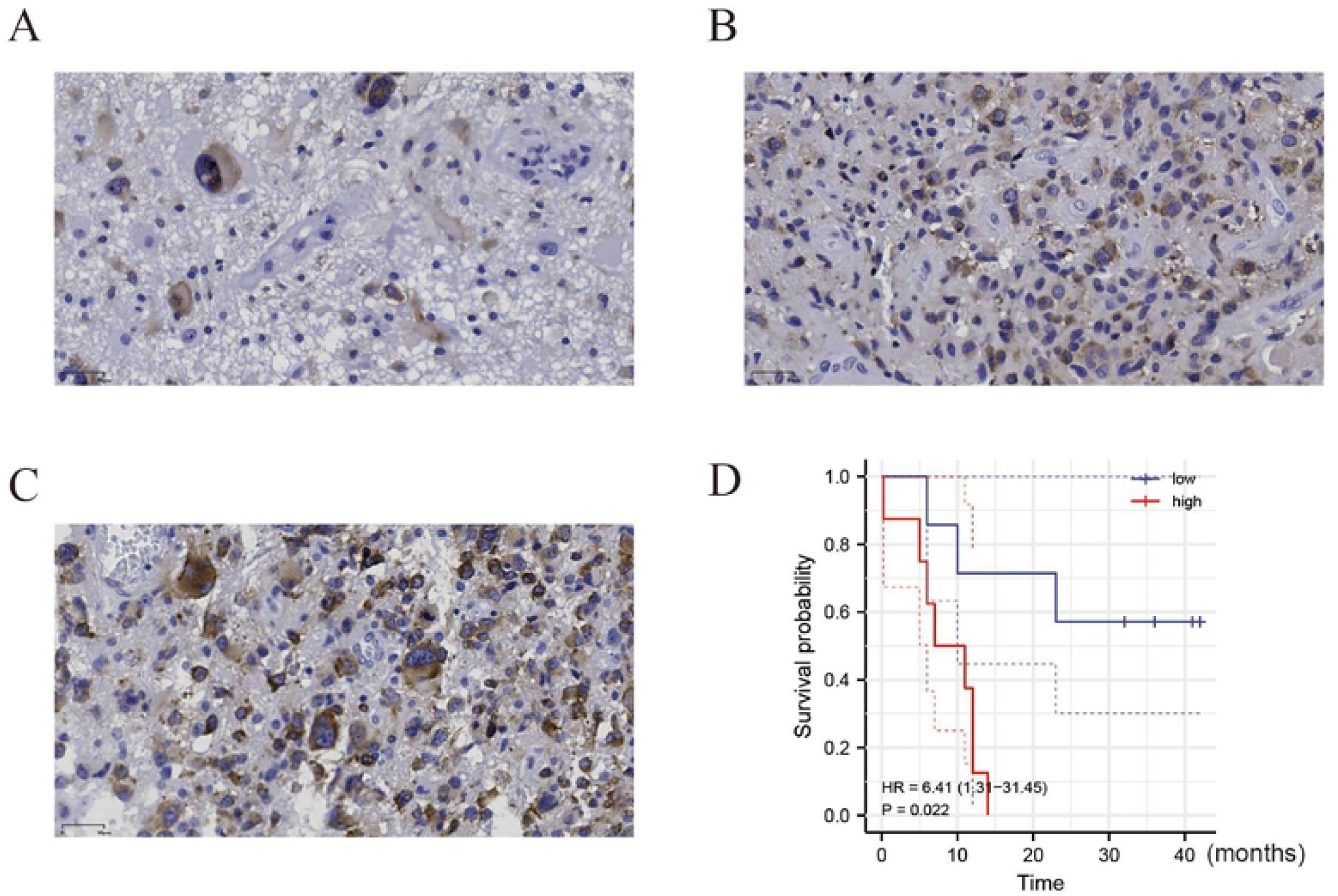
The expression of FCGRT in gliomas and its prognostic significance were analyzed by immunohistochemistry. A, B, and C showed that DDOST is weakly strongly, moderately and strongly positive in gliomas respectively. High FCGRT expression in glioma was related to poor OS (D).

### 3.3. Immuno-infiltration analysis showed that FCGRT was closely related to the tumor microenvironment

We further explored the relationship between FCGRT and the tumor microenvironment through four immune infiltration analysis methods, namely ssGSEA, ESTIMATE, EPIC, and xCell. ESTIMATE analysis showed that FCGRT was negatively correlated with tumor purity, but positively correlated with immune score, stromal score, and ESTIMATE score (Figure 3A). It was suggested that the higher the expression of FCGRT, the lower tumor purity and the higher immune score, stromal score, and ESTIMATE score. The ssGSEA showed that the higher the expression of FCGRT, the greater the number of types of immune cells were infiltrated (Figure 3A). According to the expression data, EPIC can analyze the proportions of eight types of immune cells such as B cells, CAFs, CD4+ T cells, CD8+ T cells, endothelium infiltration, endothelial cells, macrophages and NK cells. Using EPIC, we calculated the correlation between FCGRT expression and the content of eight immune cells. FCGRT was strongly positively correlated with macrophage expression (r=0.74, P< 0.0001) (Figure 3B). In order to further verify the relationship between FCGRT and macrophages, the xCell analysis method was used. The results further confirmed that FCGRT closely related to macrophages, especially M1 macrophages, with a correlation of 0.818, P< 0.001. FCGRT expression was also correlated with M2 macrophages, and the correlation coefficient was 0.518, P< 0.001(Figure 3C-E).

**Figure 3.**
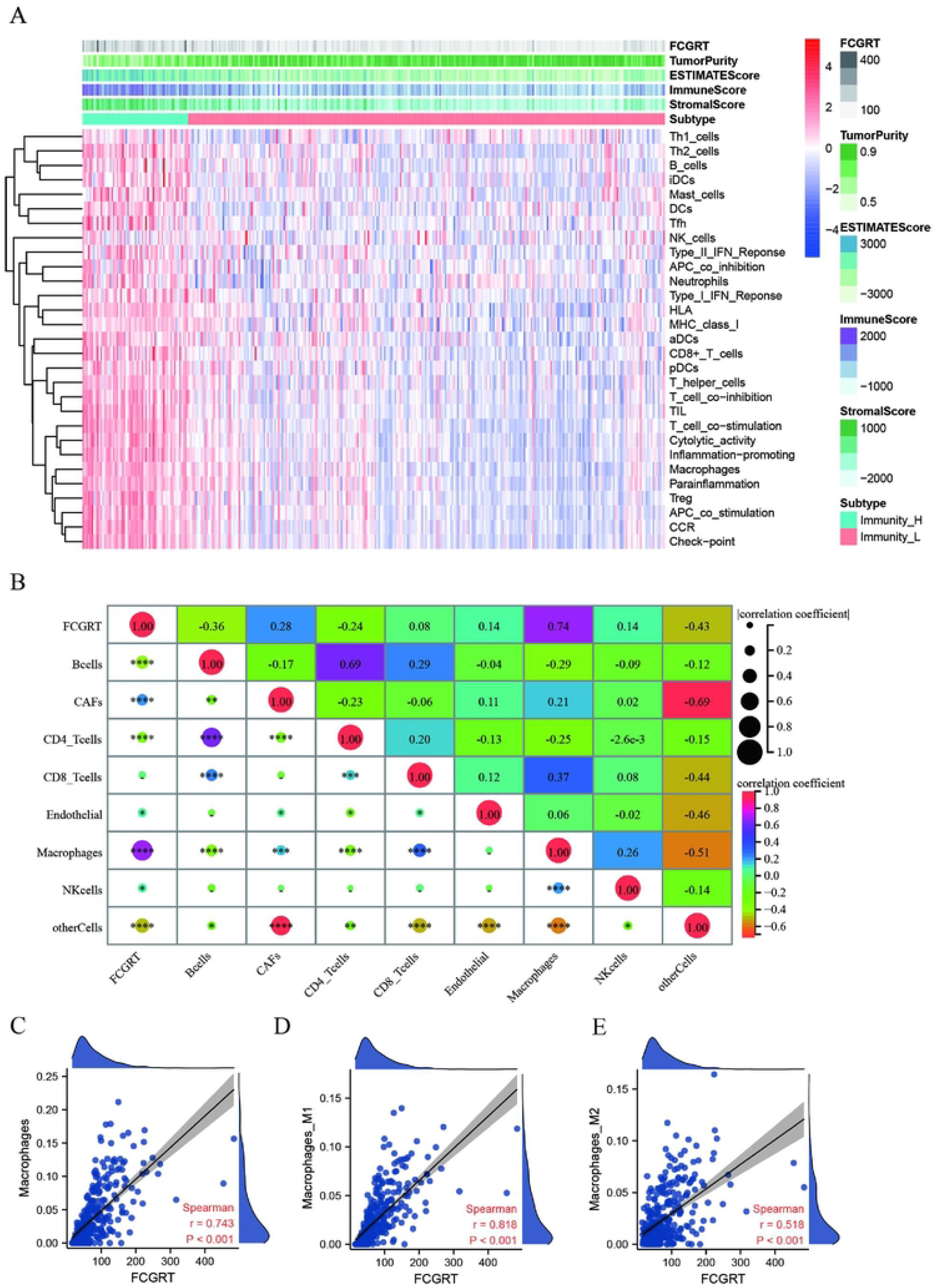
Immune infiltration patterns of low- and high-FCGRT analyzed by ssGSEA, ESTIMATE, EPIC, and xCell methods in glioma from the CGGA dataset. (A) Heatmap revealing the scores of immune cells in low and high immunities by ESTIMATE and ssGSEA. (B) Through EPIC analysis, the correlation heat map showed the correlation between FCGRT and immune related cells. (C-E) Scatter plot showed the correlation between FCGRT and Macrophages, Ml macrophages and M2 macrophages.

We obtained 109 characteristic genes of monocyte macrophages. After removing the genes from mice, the remaining 95 characteristic genes are shown in Supplementary TableA. According to NMF cluster analysis, glioma patients in the CGGA database were divided into cluster 1 (C1) and cluster 2 (C2) (Figure 4A). Survival analysis showed that results were consistent with the previous results. Patients with high expression of FCGRT had a poor prognosis (Figure 4B). By comparing the expression of FCGRT in C1 and C2, we found that FCGRT expression in C1 was significantly higher than that in C2 (Figure 4C). In previous studies, our research group found that FCGRT interacts with tumor-derived immunoglobulins. Here, we found a positive correlation between FCGRT and IGHG1 expression, which also verified our previous results (Figure 4D-E).

**Figure 4.**
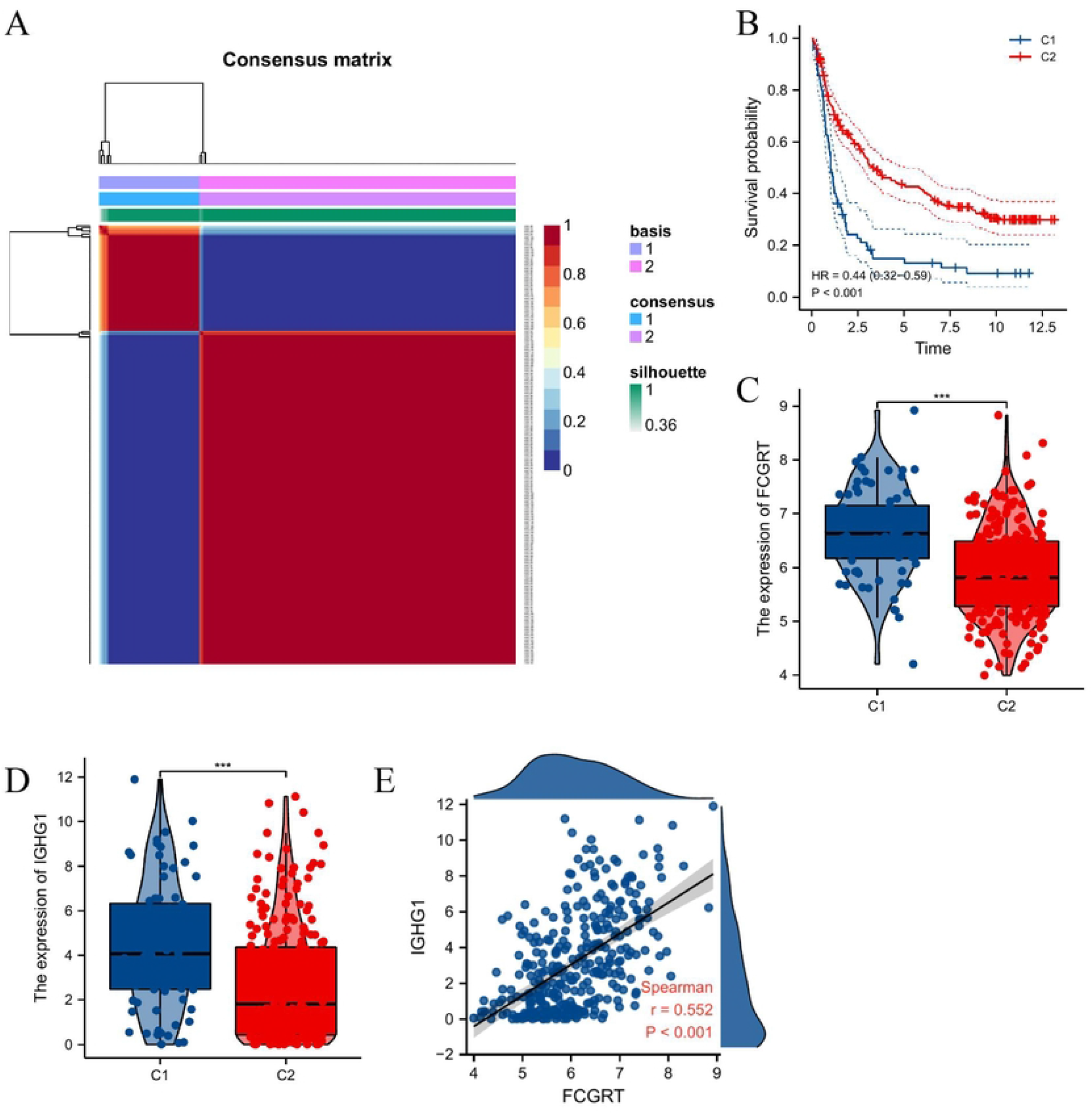
The relationship between FCGRT and monocyte macrophages related genes by NMF cluster analysis. A: According to the cophenetic value and the expression of monocyte macrophages related genes, glioma patients were divided into two clusters, Cl and C2. B: The survival curve was used to analyze the survival difference between C1 and C2 groups. C: The expression differences of FCGRT in Cl and C2 clusters were compared. D: The expression differences of IGHG1 in Cl and C2 clusters were compared. E: The scatter plot showed the correlation between FCGRT and IGHG1 expression.

### 3.4. Single cell analysis reveals that FCGRT is closely associated with macrophages and monocytes

Single cell sequencing data were downloaded from GSE117891. Data contained 6148 cells derived from 73 surgery points from 13 gliomas and 1 brain metastasis patient. The R package Seurat was used for single cell data processing and clustering. According to the characteristic gene expression of each cell, we divided these cells into five subgroups, including macrophages, astrocytes, monocytes, epithelial cells, and T cells. Supplementary Figure A3A depicts the expression landscape of FCGRT into five clusters of cells. The highest levels of FCGRT expression were found in macrophages and monocytes. Expression levels of FCGRT are shown in Supplementary Figure A3B, further confirming that FCGRT expression was highly correlated with macrophages and monocytes.

### 3.5. FCGRT downregulation attenuates proliferation, migration and invasion in human glioma cells

The above results all confirmed that FCGRT can serve as an independent prognostic factor for glioma patients being related to immune-related biological progress that regulates the glioma character phenotype. We explored the mechanism of FCGRT-mediated malignant phenotype of gliomas by GSEA analysis. We found that the function of FCGRT was enriched in apoptosis and EMT. The results of the GSEA analysis suggested that FCGRT may mediate the malignant phenotype of glioma patients by affecting the proliferation, invasion and migration of glioma cells (Figure 5A). To clarify the role of FCGRT in the context of proliferation, migration and invasion of glioma cells, we first knocked down FCGRT expression in glioma cells. Results showed that glioma cells that were transfected with FCGRT siRNA had reduced FCGRT expression (Figure 5B, C). As the inhibitory effect of siRNA-1 on FCGRT expression levels was more significant than siRNA-2, we chose siRNA-1 to investigate the mechanism(s) by which FCGRT affect the malignant progression of glioma cells in all subsequent experiments. To further confirm the role of FCGRT in proliferation of glioma cells, we examined the effects of FCGRT knockdown on proliferation in glioma cells, as shown by MTS assay and colony-forming unit assays; results showed that FCGRT significantly inhibited proliferation of U251 cells (Figure 5D–F). Moreover, a scratch wound healing assay and Boyden chamber Transwell assay showed that FCGRT knockdown clearly attenuated U251 cell migration compared with vehicle-treated cells (Figure 5G, H), and enhanced the effect of invasion of glioma cells (Figure 5I-J). To validate these observations, time-lapse imaging experiments were also performed, and similar results were obtained (Figure 5K-L). Taken together, these data demonstrate that silencing of FCGRT in U251 cells reduced the malignant progression of glioma cells.

**Figure 5.**
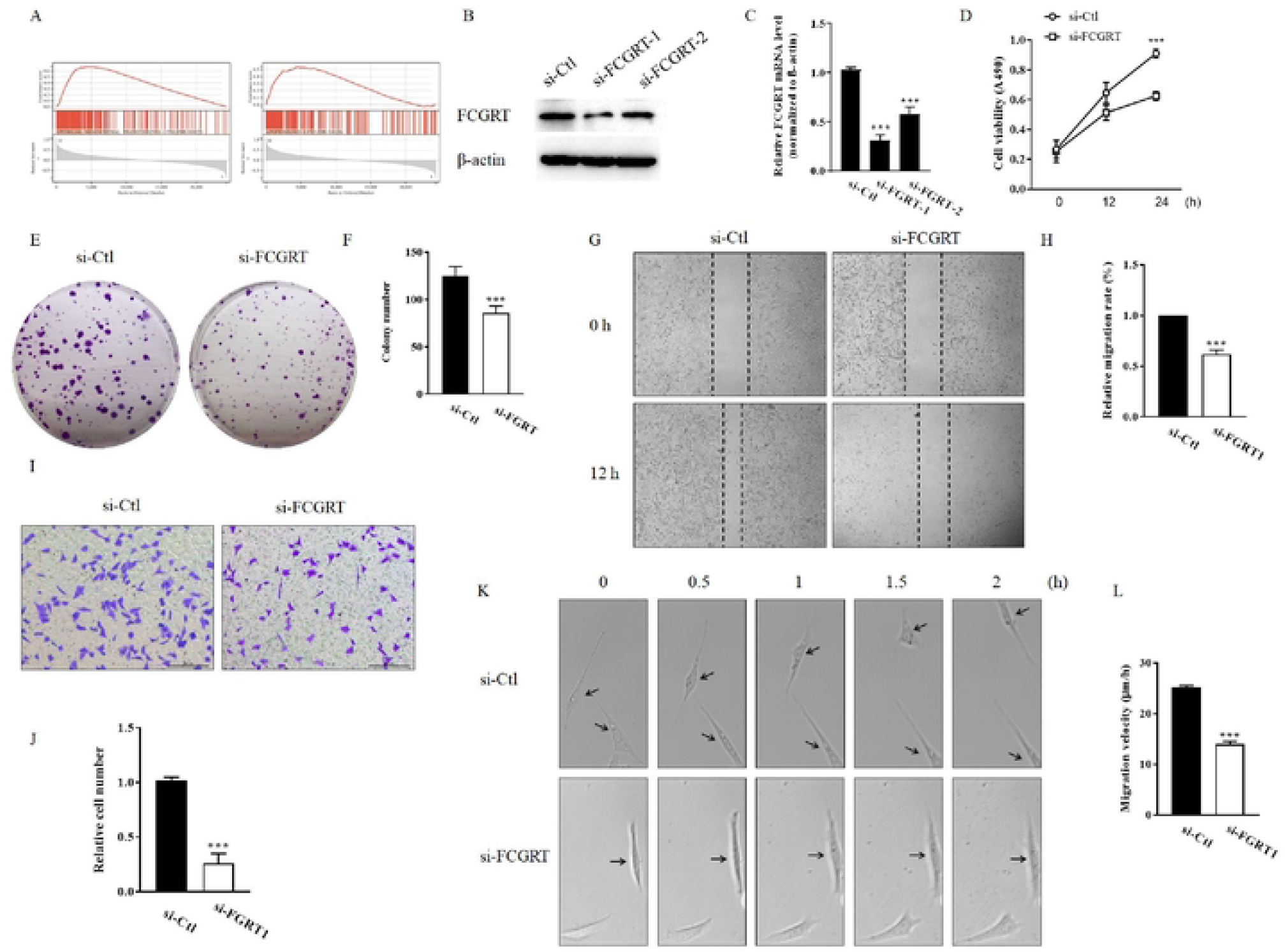
Silence of FCGRT in U25l reduced the malignant progression of glioma cells. (A) GSEA results suggest that FCGRT was associated with tumor cell proliferation, apoptosis and EMT. (B-C) U25l cells were transfected with si-Ctl, si-FCGRT-1 or si-FCGRT-2 for 24 h. The expression of FCGRT was analyzed by Western blot (B) and qRT-PCR (C). (D) U25l cells were transfected with si-Ctl or si-FCGRT for indicated times and cell viability was determined by MTS assay. (E-F) Colony formation of U25l cell was assayed and quantitated after U25l cells were transfected with siCtl or si-FCGRT. (G-H) U25l cells were treated as in (D), a scratch wound assay was performed and photomicrographed. Relative migration rate was measured by Image J. (I-J) Boyden chamber assay showing U251 cells traversed the filter to the other side and stained by crystal violet and quantifcation of cells migrated to the lower side of the membrane. Scale bars=2OO μm. (K) Migration of U25l cells transfected with si-FCGRT were traced by using time-lapse imaging. Migration distances of cells per 30 min were measured by Image J. All results are presented as the mean + SEM from 3 independent experiments (n=3). ***P<0.001 versus the corresponding control.

## 4. Discussion

Gliomas generally have a poor prognosis, especially high-grade gliomas. Currently, treatment of glioma includes surgery, chemotherapy, radiotherapy, targeted therapy and immunotherapy. However, the prognosis of glioma is still unfavorable. Therefore, it is necessary to develop new potential therapeutic targets for gliomas[18].

In recent years, increasingly more researchers have found that tumor cells can produce IgG, known as cancer-derived immunoglobulin G (CIgG)[7,19–21]. Many studies have shown that CIgG plays an important role in tumor progression. Xu found that CIgG may promote the disease progression of prostate cancer by regulating SOX2[8]. Liao et al. found that tumors with high CIgG expression of have the characteristics of tumor stem cells, with stronger spheroidizing ability and resistance to chemotherapy[22]. Our group also demonstrated for the first time that CIgG was expressed in gliomas, and patients with high CIgG expression had a worse prognosis[9]. We subsequently found that CIgG interacts with FCGRT by protein mass spectrometry. In this study, we initially explored the role of FCGRT in the progression of gliomas by bioinformatics and in vitro experiments, in order to provide a basis for future studies on the mediation of the malignant phenotype of glioma by CIgG.

FCGRT encodes a receptor that binds the Fc region of monomeric immunoglobulin G which transfers immunoglobulin G antibodies from the mother to the fetus through the placenta, and binds immunoglobulin G to protect antibodies from degradation[10]. Binding of FCGRT to IgG is performed through the FC terminal CH2-CH3 hinge region of IgG. FCGRT is widely expressed in epithelial and endothelial cells of many tissues in the body, as well as in a variety of immune cells (e.g., monocyte macrophages). Vascular endothelial cells are the main site where FCGRT protects IgG from being catabolized[23]. The scientific community has recently been intrigued by FCGRTrelated immune functions in humoral immune responses and cancer immunosurveillance. Our previous research results also found that FCGRT can bind to CIgG, but its role has not been defined.

We analyzed the effect of FCGRT on the prognosis of glioma patients. High expression levels of FCGRT indicated a poor prognosis. Some researchers have found that overexpression of FCGRT can promote tumor progression[24]. These investigators suggested that overexpression of FCGRT protects albumin from degradation in vivo, which in turn provides a sufficient source of amino acids for tumor cells to promote tumor growth. Paradoxically, some studies have reported that low expression of FCGRT can promote tumor cell growth and was associated with poor prognosis in tumor patients[25–27]; this is thought to be due to FCGRT-mediated recycling of albumin that reduces amino acid availability to fuel metabolic pathways. Furthermore, single cell and immune infiltration analysis showed that FCGRT was closely related to the immune microenvironment, especially macrophages. FCGRT is expressed on both vascular endothelial cells and circulating monocytes. These cells, through endocytosis, take up IgG from the serum intracellularly, which binds to FCGRT in acidified endosomes. Proteins that cannot bind to FCGRT are degraded in lysosomes, whereas IgG binding to FCGRT is released back into the blood circulation, prolonging its halflife[28]. This also seems to explain why our single-cell data and immune infiltration analysis showed that FCGRT was mainly expressed in monocytes and macrophages.

In vitro experiments, we observed changes in glioma cell proliferation and invasion ability by knockdown of FCGRT. This suggested that downregulation of FCGRT could inhibit the proliferation and invasion ability of glioma cells. Paradoxically, Emilie et al. found that in lung cancer, FCGRT was downregulated and was significantly correlated with poor prognosis[26]. Kristi et al. found that low expression of FCGRT was related to a poor prognosis in colorectal cancer patients. The main reason for this is that FCGRT mediates the intestinal mucosal immune response. In the absence of FCGRT, the mucosal anti-tumor immune response cannot be effectively activated[29]. In gliomas, we found that glioma cells with high expression of FCGRT had stronger proliferation and invasion ability. This result was contrary to the conclusions of other researchers. There are two possible reasons for this. First, there is a huge difference between the intracranial microenvironment and the body tumor microenvironment; second: other scholars’ studies are in vivo experiments, and may pay more attention to FCGRT expressed by macrophages or endothelial cells, while our research was mainly limited to in vitro experiments, which paid more attention to FCGRT expressed by tumor cells.

This study is a follow-up to the earlier study of our research team. We previously found that CIgG mediated the poor prognosis of glioma patients. Mass spectrometric experiments showed that CIgG could bind to FCGRT. Furthermore, in this study, we analyzed the significance and possible mechanism of FCGRT in gliomas. However, there are still some deficiencies in this study. In future studies, we will try to observe the effect of FCGRT on glioma cells in vivo.

## Author Contributions

Guohui Wang, Zheng Wang, Tingting Zhang: Performed most of the experiments. Hongyao Ge, Jie Pan, Wangyang Yu, Tianfang Yan: Performed part of the experiments, Prepared the manuscript. Gaoshan Yang and Wei Jiang: Designed the study, Supervised the research, Prepared the manuscript writing, Revised the manuscript.

## Institutional Review Board Statement

The study was conducted in accordance with the Declaration of Helsinki, and approved by the Institutional Animal Care and Use Committee of Hebei University of Chinese Medicine au-thorized all processes (no. DWLL2020018).

## Data Availability Statement

RNA sequencing data and corresponding clinical information of TCGA LGG and GBM datasets were downloaded from TCGA data portal (https://portal.gdc.cancer.gov/). Gene expression profiling and corresponding clinical features of gliomas were obtained from the CGGA database (http://www.cgga.org.cn/). The dataset GSE117891 for single cell analysis was downloaded from the GEO database (https://www.ncbi.nlm.nih.gov/geo).

